# STABIX: Summary statistic-based GWAS indexing and compression

**DOI:** 10.1101/2024.11.15.623812

**Authors:** Kristen Schneider, Simon Walker, Chris Gignoux, Ryan Layer

## Abstract

Genome-Wide Association Studies (GWAS) are widely used to investigate the role of genetics in disease traits, but the resulting file sizes from these studies are large, posing barriers to efficient storage, sharing, and querying. This issue is especially important for biobanks like the UK Biobank that publish GWAS for thousands of traits, increasing the volume of data that must be effectively managed. Current compression and query methods reduce file sizes and allow for quick genomic position-based queries but do not provide utility for quickly finding loci based on their summary statistics. For example, finding all SNVs in a particular p-value range would require decompressing and scanning the whole file. We propose a new tool, STABIX, which introduces summary-statistic-based queries and improves upon the standard bgzip compression and tabix query tool in both compression ratio and decompression speed. When applied to ten GWAS files from PanUKBB, STABIX created smaller compressed data and indices than tabix for all files, where bgzip and tbi files were an average of 1.2 times the size of STABIX compressed files and indexes. In the same ten files, STABIX per gene decompression was, on average 7x faster than tabix per gene decompression, and achieved faster per gene decompression times for over 99% of nearly 20,000 genes.

## INTRODUCTION

### Genome-wide association studies

(GWAS) use statistical approaches to identify connections between genotypes and phenotypes by looking across control and disease populations (e.g. heart disease, type II diabetes, auto-immune and metabolic disorders, etc.) to identify single-nucleotide variants (SNVs) that are likely associated with certain disease traits^1,2^. Some of the earliest GWAS identified independent SNV association signals in bipolar disorder, coronary artery disease, Crohn’s disease, rheumatoid arthritis, and other common diseases^3^. These early findings inspired hundreds of other studies which aimed to look more closely at single diseases and to explore disease patterns in diverse populations. Despite some of their early critiques (e.g. unclear biological relevance, flawed assumptions, and spurious results), GWAS has maintained its popularity for the last 15 years as researchers continue to improve analysis methods and publish new GWAS-based discoveries^4–6^.

The recent and ongoing surge in biobanks (UK Biobank^7^, deCODE genetics, Biobank Japan^8^, etc.) has played a key role supporting the popularity of GWAS^9^. The larger sample sizes in these biobanks provide greater statistical power to discover new associations^10^. GWAS for height, blood pressure, and smoking initiation, for example, have made history with sample sizes of over one million. In 2015, the National Human Genome Research Institute-European Bioinformatics Institute (NHGRI-EBI) redesigned its GWAS Catalog database to support the influx of GWAS, as well as support a wider range of information included with the studies (e.g. ancestry and recruitment information)^2,11^. As of August 2024, this catalog contains nearly 67,000 top associations and more than 90,000 full summary statistics. Additionally, the most updated version of Pan UK Biobank (PanUKBB, v0.4) includes a multi-ancestry set of over 7,000 GWAS for a half million samples from 6 continental ancestry groups^12^.

Beyond the fundamental exploration of trait-disease associations, large-scale GWAS can help derive predictions about patients’ disease risks, termed polygenic risk scores (PRS)^13,14^. These measurements estimate a patient’s risk for certain diseases based on the unique variants in that patient’s genome. When applied appropriately PRS can raise awareness of diseases before symptoms arise, inform decisions that can help slow disease progression, and help identify targets for drug development^15^. The power of large sample sizes allows for a deeper characterization of a patient’s medical profile and advances our understanding of the genetic landscape of complex diseases. Even still, several untapped applications for GWAS lie just beneath the surface, such as the inclusion of diverse populations, alternative inheritance models, and other biological and environmental metrics^16^. One factor that limits this potential is data size. Summary statistics alone can require hundreds of terabytes of space in a single biobank, making storage, sharing, and downstream investigation challenging.

With current technologies, GWAS summary statistics for a single trait require between 3 and 11GB of storage for bgzipped and plain text data, respectively. For over 7,000 traits in the current PanUKBB, this can amount to over 10TB of compressed data. This large and growing collection of GWAS ensures continued interest in sharing summary statistics across research and clinical communities and requires new computational methods for efficient storage and computation.

### Format-specific compression methods

Two of the most ubiquitous methods that improve the accessibility of GWAS data use compressed (bgzip) and indexed (Tabix) data to reduce the storage burden and time required to retrieve individual records^17^. Bgzip is similar to gzip but introduces block-based compression, allowing for efficient regional data decompression. Tabix works alongside bgzip to create a genomic position index of the compressed blocks, enabling quick position-based data retrieval. While the standard B-tree index froml SQL databases is a reasonable approach to perform these queries, tabix is preferred to SQL approaches as it directly works with popular file formats, uniquely works on compressed data files, and is faster than SQL databases at for big data[https://github.com/samtools/tabix/blob/master/tabix.1]. While bgzip and tabix facilitate storing and retrieving GWAS summary statistic data (e.g. BED file formats), their compression is limited to a single compression scheme (i.e. codec), and the access pattern is restricted to genomic position-based queries.

To strengthen the efficient investigation of GWAS data, we introduce STABIX, a compress and index method that improves upon bgzip compression ratios with a ensemble codec compression approach, and improves upon tabix queries with the addition of a summary-statistic-based index. STABIX adds column compression to bgzip’s block-based structure, allowing multiple codecs to be used for the different data types, resulting in a more effective compression strategy. Additionally, STABIX extends the tabix position-based index to include a summary-statistic index, supporting a more comprehensive range of GWAS data exploration strategies. To accomplish this, first, STABIX creates blocks of rows of fixed or variable (see Methods) length and compresses individual columns with codecs corresponding to their data type (**Figure 1A**). Concurrent to compression, STABIX generates a genomic index which stores block, genomic, and file information necessary to reconstruct individual blocks. Optionally, a second index can be created for some column of interest in the original GWAS file (e.g. p-value) which will permit efficient search which satisfies some constraint (e.g. p-value <= 5e-8) (**Figure 1B**). Finally, STABIX’s query step allows users to query with both genomic and statistical contraint (**Figure 1C**). The STABIX software is openly available at https://github.com/kristen-schneider/stabix.

**Figure 1.**
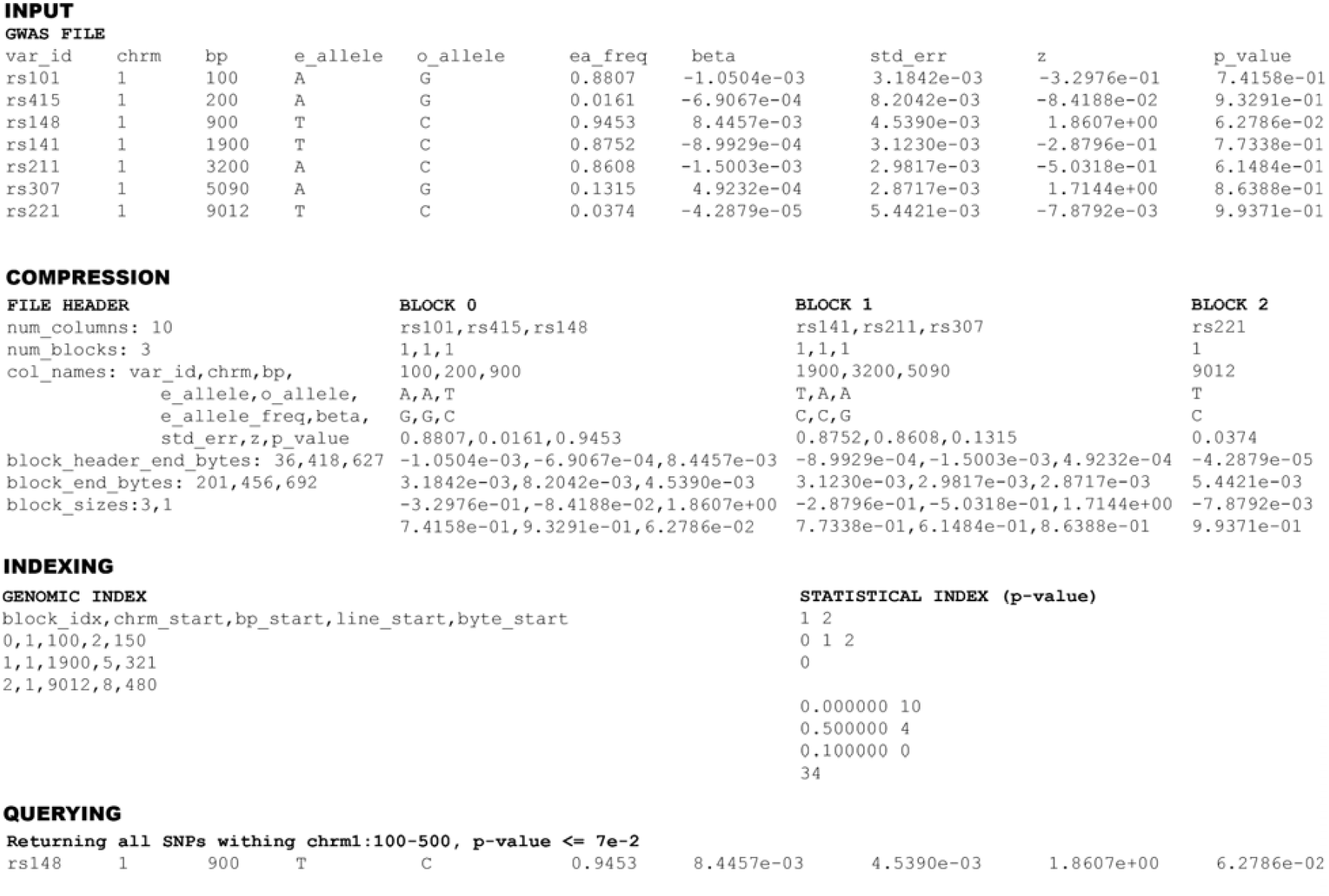
– STABIX workflow. STABIX takes a single GWAS file and configuration file as input. STABIX compression separates GWAS data into blocks, and generates a column-centric data structure for column-based compression. STABIX generates a genomics index (genomic positions) and a statistical index (statistical bins) specified by the configuration file. STABIX returns records which match a query specified by the configuration file.

## RESULTS

### Finding Genome-Wide Significant SNVs

STABIX’s unique summary-statistic index offers efficient access to SNVs based on the strength of their association with the target trait. By only inflating blocks that contain a SNV with an association above the user-defined threshold, STABIX avoids a substantial amount of wasted work. Since tabix index stores no information about the underlying summary statistic data, it must inflate all blocks first before passing the output to a second method that parses and tests the association values. To measure this improvement, we selected ten PanUKBB GWAS files and queried 19,181 protein-coding genes for SNVs with p-values at or below 5e-08 using default parameters. STABIX was faster than its equivalent tabix workflow for over 99% of gene queries (**Figure 2A**; see **Supplementary Table 1** for codec ensemble descriptions). As expected, the speedups depend on whether the target genes had significant SNVs (**Figure 2C**). For the 88.6% of genes that contain zero significant SNVs, STABIX was 7.7X faster than TABIX. For the minority of genes with significant SNVs, STABIX was 1.75X faster. We run the same experiment for 100 PanUKBB GWAS files with *continuous* traits and report that STABIX is faster than its equivalent tabix workflow for 99.15% of gene queries with an average speedup of 5.41X. STABIX was faster than tabix for 93.31% of genes with significant SNVs and reports an average speedup of 1.70x over tabix (see **Supplementary Figure 1**).

**Figure 2.**
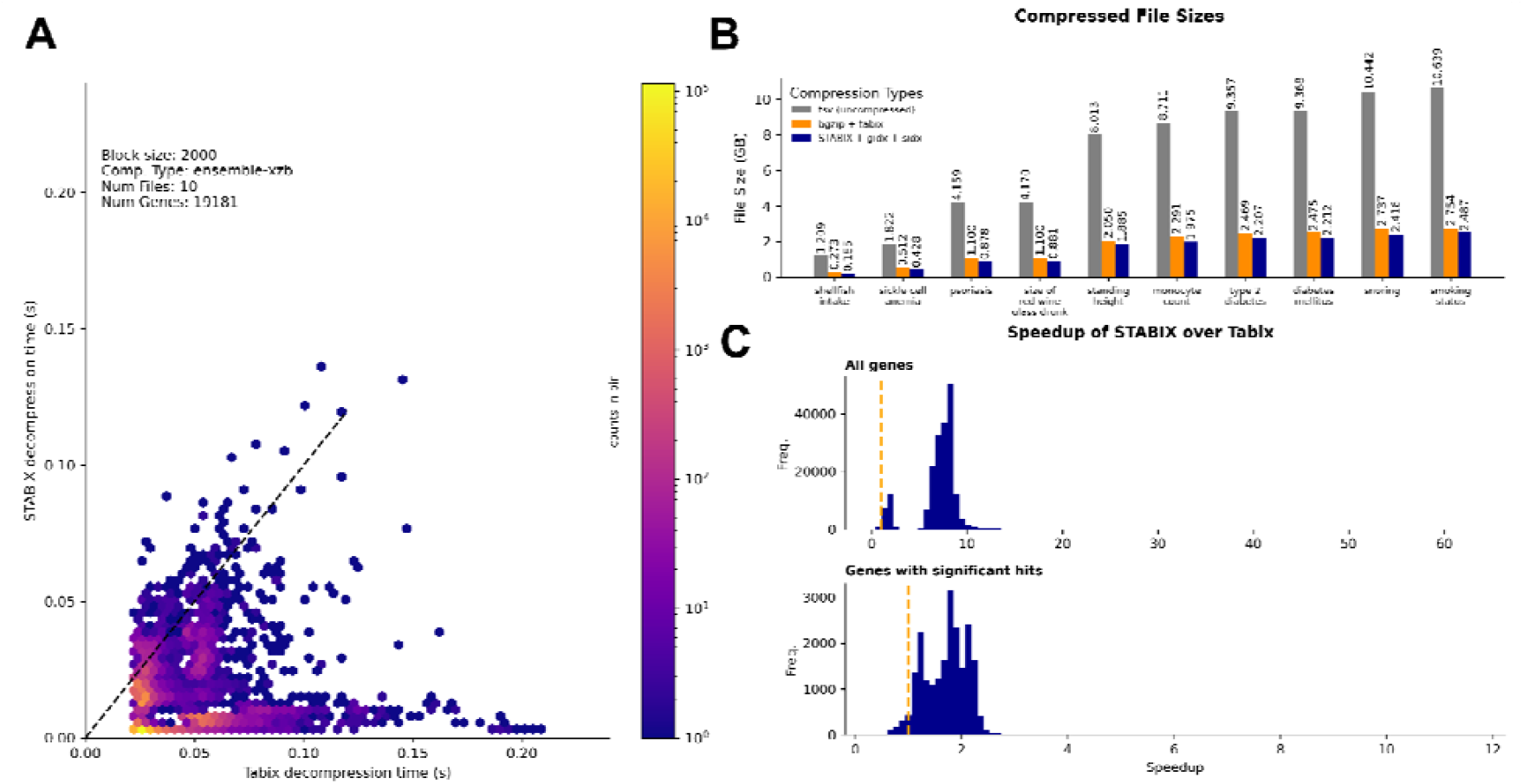
STABIX vs tabix performance for 10 PanUKBB GWAS per-phenotype files. **A** Significant SNV query speed by gene for Tabix and STABIX with hexagonal binning. **B** Uncompressed, bgzip/tabix compressed, and STABIX compressed file sizes. Bgzip/tabix compressed file sizes include the bgzip compressed file (bgz) and tabix index file (tbi). STABIX file sizes include the STABIX compressed file, the genomic index (gidx) and the statistical index (sidx) files. **C** Speedup by gene for significance SNV queries for STABIX over tabix + filtering step for all genes (top) and only genes with significant hits (bottom). A vertical dashed line (orange) is drawn at x=1 to separate where tabix wins (left of line) and STABIX wins (right of line). All figures report data for 10 per-phenotype files from PanUKBB, STABIX compression using default block size 2000 and codec ensemble xzb.

To better understand which features of each gene impacted the STABIX and tabix query time, we investigate various components of the query. **Figure 3A** demonstrates that while there is a slight increase in decompression speed for both STABIX and Tabix queries as the uncompressed data grows larger, file size is not a limiting factor for decompression time, nor an advantage to either method. **Figure 3B** shows that the number of STABIX blocks also does not account for the difference in STABIX and tabix decompression times. While STABIX and tabix design blocks with different approaches, for both, using more blocks typically aligns with slower decompression times. **Figure 3C** shows that STABIX is significantly faster than tabix when there are fewer significant p-value hits found in a block. This is because STABIX avoids unnecessary decompression when there are no p-value hits found. Likewise, **Figure 3D** shows that STABIX offers significant speedup over tabix in cases where there are few significant SNVs to return for a genomic query. To explore this phenomenon further, we select a highly heritable and polygenic trait (standing height^18^) and plot comparisons for genes with no hits, p-value hits, and p-value hits within a genomic query in Supplementary **Figure3A,B** and **C** respectively.

**Figure 3.**
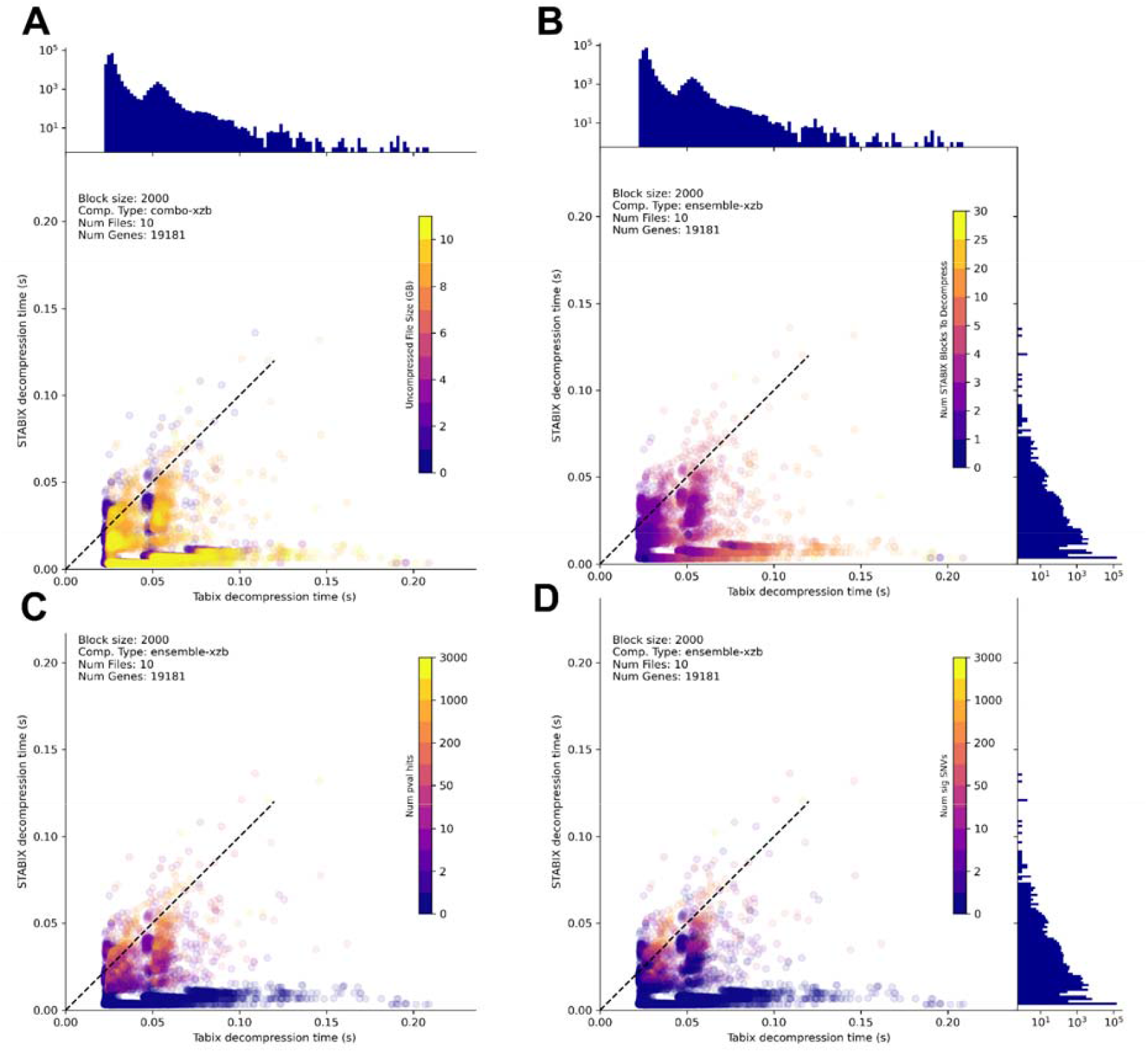
STABIX vs tabix decompression speed by gene with coloring by query information. Decompression speed by gene colored by **A** uncompressed file size, **B** number of blocks decompressed with STABIX **C** number of p-value hits found during search **D** number of significant SNVs returned in query. The frequency of genes plotted at each y-location is plotted in a log-axis histogram to the right of each row. Histograms for STABIX and tabix decompression times are shown on the right and top of grids, respectively.

### Reducing the GWAS file storage burden

In addition to enabling the indexing of statistical columns of interest, the STABIX column-based strategy provides a mechanism to enhance compression beyond what bgzip offers by employing additional codecs specifically optimized for various data types. This approach allows for more efficient storage and retrieval, as each column is compressed using methods best suited to its particular characteristics. On average, STABIX decreased file sizes by an additional 4% compared to bgzip (**Figure 2B, Table 1**). To put that into perspective, if the PanUKBB uncompressed GWAS files required 77 TB, then bgzip would reduce the burden to about 20TB, and STABIX would reduce it to 17TB. That 3TB savings would come in addition to the retrieval speedups discussed above.

**Table 1.** Performance of STABIX. Performance measurements for ten per-phenotype files from PanUKBB. STABIX compression using default parameters. A win is awarded when a method’s query time per gene is faster than the other, and percent (%) win is the average number of wins over 19,181 genes in the ten files. Compression ratios include index files when available.

| trait | shellfish intake | sickle cell anemia | psoriasis | size of red wine glass drunk | standing height | monocyte count | type 2 diabetes | diabetes mellitus | snoring | smoking status |
| --- | --- | --- | --- | --- | --- | --- | --- | --- | --- | --- |
| no. sig genes | 977 | 42 | 143 | 0 | 5776 | 1803 | 141 | 156 | 59 | 168 |
| no. sig SNVs | 2189 | 392 | 2232 | 0 | 165522 | 41759 | 3514 | 4039 | 2120 | 5004 |
| avg. STABIX speedup over tabix | 5.095 | 6.549 | 7.1693 | 7.1773 | 4.2627 | 6.7368 | 8.3546 | 8.1861 | 8.4399 | 8.2152 |
| % STABIX wins | 98.4412 | 99.9791 | 99.9583 | 100.0 | 99.2649 | 98.4777 | 99.9009 | 99.8853 | 99.9583 | 99.9531 |
| uncompressed (GB) | 1.2091 | 1.8225 | 4.1587 | 4.17 | 8.0126 | 8.7106 | 9.3567 | 9.3676 | 10.4422 | 10.6391 |
| bgzip (GB) | 0.2715 | 0.5107 | 1.0985 | 1.0982 | 2.0483 | 2.2893 | 2.4672 | 2.4732 | 2.7342 | 2.7522 |
| tabix index (MB) | 1.6885 | 1.7465 | 1.9274 | 1.9279 | 2.1941 | 2.2362 | 2.2698 | 2.2704 | 2.3161 | 2.3221 |
| STABIX (GB) | 0.1838 | 0.4277 | 0.8771 | 0.8804 | 1.8838 | 1.9746 | 2.2061 | 2.2118 | 2.4149 | 2.4858 |
| STABIX genomic index (MB) | 0.506 | 0.511 | 0.513 | 0.513 | 0.5207 | 0.5211 | 0.522 | 0.522 | 0.5226 | 0.5228 |
| STABIX statistical index (MB) | 0.1981 | 0.1534 | 0.1553 | 0.1537 | 0.2522 | 0.2024 | 0.1605 | 0.1604 | 0.1613 | 0.1698 |
| STABIX:raw text compression ratio | 6.5534 | 4.2542 | 4.7379 | 4.7328 | 4.2516 | 4.4097 | 4.2401 | 4.234 | 4.3229 | 4.2787 |
| STABIX:bgzip+tabix compression ratio | 1.4805 | 1.1963 | 1.2536 | 1.2486 | 1.088 | 1.1601 | 1.1191 | 1.1189 | 1.1329 | 1.1078 |

### Selecting default parameters for STABIX

STABIX splits data into blocks and then performs compression by column, allowing for different columns (i.e. different data types) to be compressed with different codecs. We evaluate a set of six popular codecs on three data types (e.g. integer, floating point, and string data). Selecting a single file (GWAS trait: shellfish intake), we perform STABIX compression, indexing, and decompression for each codec ensemble described in Supplementary Table 1 and block sizes 1000, 2000, 5000, 10000, and 1 centimorgan (cM). We query the set of protein-coding genes, and record decompression times and compressed data sizes for each column, and show results in **Figure 4**.

**Figure 4.**
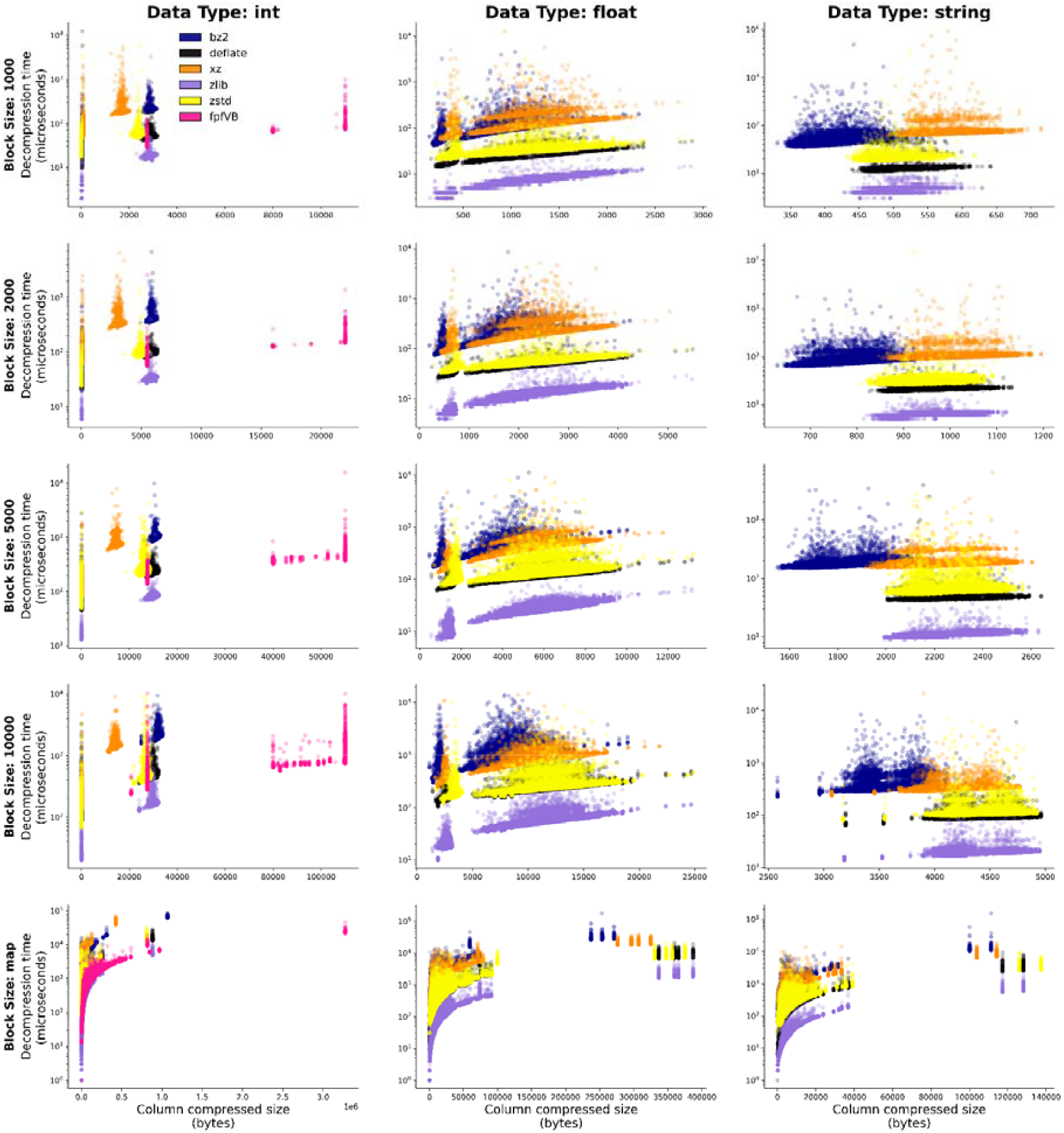
Codec performance by data type. Decompression time (log scale) vs. compressed size of a single column. Block sizes 1000, 2000, 5000, 10000, and cM-based blocks are plotted in descending rows, respectively. Column data types integer, float, and string are plotted in columns left to right, respectively. We note that the fpfVB codec only functions on integer data.

Noting patterns of quickest decompression and smallest size of compressed data, we select two ensemble codec configurations with which to run complete STABIX compression, indexing, and decompression for the same file (GWAS trait=shellfish intake) and show a subset of results from these experiments in **Figure 5**. We include all 40 combinations of block size and codec ensembles from this experiment in **Supplementary Figure 2**.

**Figure 5.**
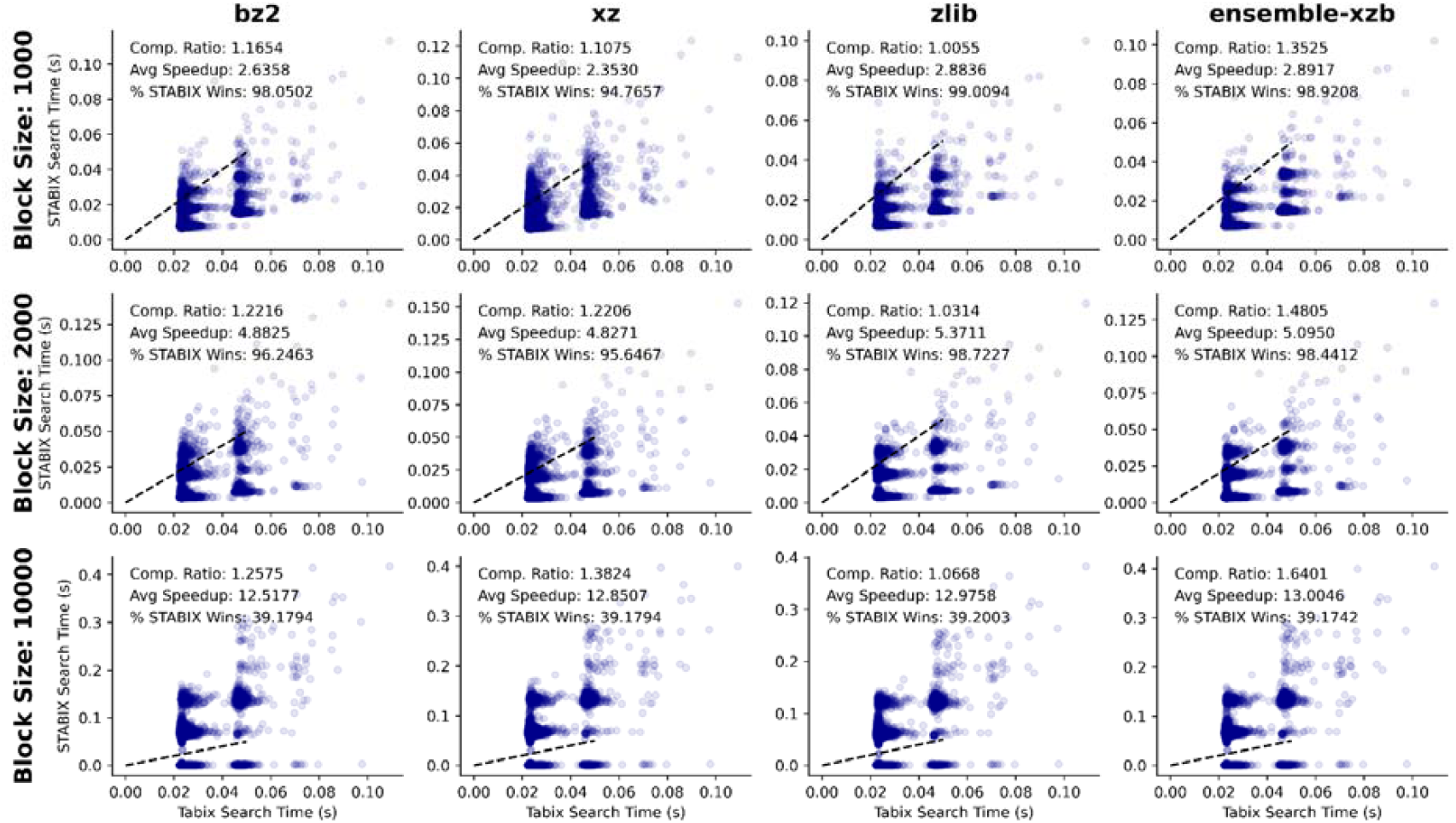
Performance of varying block sizes and codec ensembles for a single file. STABIX vs. tabix gene-based query times for PanUKBB GWAS file trait: shellfish intake. Times are shown for four codec configurations from left to right: bz2, xz, zlib, and ensemble-xzb; and across three block sizes from top to bottom: 1000, 2000 and 10000. Bgzip + tabix : STABIX + genomic index + statistical index compression ratios, STABIX average speedup, and % STABIX wins are reported for each of the twelve configurations.

From these results, we report the overall best performing configurations for compression ratio, average speedup, and percent (%) queries with STABIX speedup over tabix (% STABIX wins) in **Table 2**. In effort to maximize compression ratio, average speedup, and % STABIX wins, we select block size 2000 and codec ensemble ensemble-xzb (int=xz, floating point=zlib, string = bz2) as the default configuration. Alongside these observations we note the following three things: first, ensemble-xbb (int=xz, floating point=bz2, string=bz2) has comparable results depending on file size (see **Supplementary Table 2**). Second, while block size 10000 and codec ensemble-xzb wins for both compression ratio and average speedup, it performs poorly for % STABIX wins (where tabix beats STABIX for more genes). In this case, STABIX is able to achieve a very high speedup for a few genes, which makes STABIX faster than tabix on average, but for the majority of genes, tabix is faster. Finally, we recall that a user has the option to select their own block sizes and codecs depending on their data and desired result.

**Table 2.** Best performing codec ensemble and default parameters. For a single file, the best performance for compression ratio, average STABIX speedup over tabix, and % STABIX wins are recorded. Additionally, we include the performance of the default configuration for comparison. The winning values are highlighted in grey for each measurement, and the default parameters are additionally highlighted in grey for comparison.

|  | Codec Ensemble | Block size | Compression Ratio | Average Speedup | % STABIX wins |
| --- | --- | --- | --- | --- | --- |
| **Best compression ratio** | ensemble-xzb | 10000 | 1.6401 | 13.0046 | 39.1742 |
| **Best avg speedup** | ensemble-xzb | 10000 | 1.6401 | 13.0046 | 39.1742 |
| **Best % STABIX wins** | zlib | 1000 | 1.0055 | 2.8836 | 99.0094 |
| **Default Parameters** | ensemble-xzb | 2000 | 1.4805 | 5.0950 | 98.4412 |

## METHODS

In this section we first describe the input files necessary to complete a STABIX. Second, we detail each of the three steps which comprise the STABIX’s compression and indexing procedures. Third, we discuss a STABIX query, including what constitutes a query, and how STABIX achieves its output. Finally, we will describe methods for reproducing experiments which are reported in our results section.

### STABIX Input Files

**GWAS file,** standard bed file format (sorted by genomic position, tab delimited, includes chrm, bp, etc.). For our experimentation and results we used 10 per-phenotype files from the PanUKBB. See data and code availability below for link to PanUKBB data.

**Configuration file,** specifies STABIX options including path to GWAS file, block size, codec options, and query information. Example’s provided at: https://github.com/kristen-schneider/stabix/blob/main/config_files/test_config.yml.

### STABIX Compression and Indexing

#### The STABIX Header

STABIX’s custom header stores necessary information required to fully reconstruct (i.e. decompress) the original file and return queries. This header includes: number of columns in the original GWAS file, number of blocks created during compression, a list of the column headers in the original GWAS file, a list of block header end bytes, a list of block end bytes, and a list of block sizes (i.e. last block might be different than specified block size, or all block sizes are different if a map file is used to create blocks.) Some of the header elements can be obtained without moving through any compression steps (e.g. number of columns in the original GWAS file). Others cannot be included in the header until after data compression is complete (e.g. block end bytes). Generating and writing the header is not completed until compression is completed.

#### Block-based compression

In many cases, it is not necessary to decompress a complete file (or other representation of data) in order to accomplish a task or answer a question. For example, with GWAS, it is common that a researcher might only be interested in looking through a particular region (i.e. gene or chromosome), or that they might only be interested in data that falls at or below some p-value. Block-based compression provides the benefit of decompressing only part of the data to investigate a query, over the need to decompress a full file. This feature can decrease decompression time when queries are small, or span disjoint regions.

To achieve block-based compression for meaningful queries, a genomic index is created to store genomic information (e.g. block 1 = chromosome 1, base pairs 100 - 1300), byte-locations, and other information necessary to decompress a single block. Additionally, a binning index is created to store information about what blocks contain what kinds of data (e.g. blocks 1, 8, and 13 contain records with p–values at or below 5e-8.). More about how these indexes are created can be read about in the genomic and statistical index sections below.

The practicality of block-based compression depends on the size of the blocks as it relates to the queries being performed. Blocks with only a few lines might demonstrate quicker decompression times, but will have a larger overhead for full compressed file size (i.e. a greater number of blocks requires a more lengthy header to account for individual blocks). Furthermore, if the blocks are too small compared to the size of a typical query, the number of blocks which would need to be decompressed could end up reversing the benefit of quick decompression times. On the other hand, block sizes with a large number of lines might experience a greater decompression ratio (though not necessarily), but the decompression time starts to slow^19^.

To achieve block-based compression, STABIX parses the input GWAS file into a set of blocks whose size is determined by the user in our configuration file. STABIX’s default block size is fixed at 2,000. Optionally, the user can set the block size to point to a genomic map file (examples provided in our GitHub repository). In this case, block sizes are set to be 1cM in length, and contain a variable number of records.

#### Column-specific compression

Within each block, records (i.e. rows) are further split into columns. All data from a single column in a single block is stored together in a list. Each column is then compressed independently, with the codec specified in the config file (codec specified by data type). After a column is compressed, the end byte of that column is recorded in the block’s header. Finally, the block header is compressed with the zlib codec. A compressed block constitutes a compressed block header, and a list of compressed columns whose length is the same as the number of columns in the file.

#### Writing compressed data

After each block has been independently compressed we record the end bytes of the compressed block. The end bytes of each compressed block header and the end bytes of the compressed block are stored in the overall compressed file header as described above. The file header is compressed also with the xz codec; and the size of this compressed data is stored in the first four bytes of the compressed file. After the first four bytes, the compressed file header is written, then the first compressed header of the first block, then the first compressed block, then the second, and so forth.

#### Genomic Index

The genomic index is created to make searching for genomic coordinates (e.g. return all records from chromosome 1 base pairs 12345-56789) more efficient. Because the input GWAS files must be provided in sorted order (i.e. by increasing chromosome and base pair coordinates), the genomic index is created concurrently with the compression step. As a block is being compressed, the genomic index records the block’s index, the chromosome at the start of the block, the base pair at the start of the block, the line number at the start of the block, and the block’s byte offset. Every block is compressed independently, and the genomic index is written after the last step of the compression process.

#### Statistical Index

The floating point-based, binning index is created to make searching within a threshold of some query statistics (e.g. all records with p-value at or below 5e-8) more efficient. In this case, the input file is not sorted by this statistic, so the binning index is not created concurrently with compression. First, the user specifies bin boundaries and a query threshold in the configuration file. Because the UKBB p-values are stored as -log_10(p-value) to avoid underflow, STABIX creates 3 default bins: less than 0.3, between 0.3 and 4.3, and greater than 7.29; and a default query threshold greater than or equal to 7.3 (-log_10(5e-08) ∼ 7.3). While our results and defaults demonstrate utility for p-value statistical indexing, STABIX can create a similar index for any statistical column of interest, with any user-defined bins, and query thresholds (e.g. less rigid p-values). At index time, each bin is assigned all record IDs that fall within the bin’s range based on the record’s corresponding value in the specified column (e.g. p-value). All records are guaranteed a bin ID unless the column-of-interest value is unavailable, regardless of the bin thresholds. This operation is linear in the number of records. The binning index is serialized uncompressed. At query time, bins that overlap the query threshold are identified conservatively and their included record IDs are returned in aggregate. Extraneous record IDs can occur due to the coarse nature of the binning process and are filtered out before being presented to the user.

#### The STABIX query

STABIX first uses the statistical index to determine a set of blocks which contain records that satisfy the statistical threshold (e.g. p-value <= 5e-08). Then, for each query (e.g. gene), STABIX uses the genomic index to determine which blocks fall within the query boundaries (e.g chrm1:12345-67890). During this step, STABIX searches through a map of base pair positions for each chromosome in log(n) time to quickly return matching blocks. STABIX takes the intersection of these two sets (statistical hits ∩ genomic hits) to determine a final set of blocks to decompress. For each block in this set, STABIX first decompresses only the column specified by the statistical index (e.g. p-values) and identifies which specific indexes in this column satisfy the statistical threshold (STABIX has already determined that this block contains at least one record which satisfies the statistical threshold, and now needs to determine the specific set of records). Once these indices are determined, STABIX decompresses only the chromosome and base pair columns to determine whether or not this statistical hit falls within the gene (STABIX has already determined that the gene is contained in this block, but must determine if each record is within the specific boundaries of the gene). If at any point during these checks, a block no longer contains a hit (e.g. a p-value hit is just outside the boundary of the query), STABIX moves to the next block. Next, STABIX decompresses the appropriate set of blocks back into the column-centric readable data that was generated in the column-specific compression from above, allocates space for only the set of records which meet both criteria (statistical and genomic hit), and transposes only those matrix locations to return.

#### Experimentation

For a single file, we provide a bed file of 19,181 protein-coding genes and a p-value threshold <= 5e-08 (7.3 after -log_10(p-value) transformation by PanUKBB). We write a Python script with the pysam library to perform tabix queries for each gene in the bed file adding a statistical check to compute p-value criteria. We perform STABIX compression, indexing, and decompression according to the appropriate configuration; and check that Tabix and STABIX output match. In the case of four genes across ten files (TNIP2, SHPRH, MCCD1, and BNIP3L), STABIX reports 1 more record than tabix, where STABIX is inclusive with a base pair boundary and tabix is not. We compute STABIX average speedup over tabix by computing [tabix_time / STABIX_time] for each gene, and taking the average over these measurements for a single file.

## DISCUSSION

STABIX is a novel compression and indexing framework that builds upon the widely used combination of bgzip and tabix by incorporating data-dependent compression codecs and built-in filtering summary statistics for statistical-based queries. Both of these features allow for STABIX to improve upon decompression speed (**Figure 2A**) and compression ratio (**Figure 2B**) as compared to bgzip and tabix. With this in mind, we emphasize that STABIX’s addition of the statistical index provides its most significant advantage over tabix as it allows for sophisticated searching beyond block-based decompression.

Column-based compression offers customized compression for different data types, optimizing compressed file sizes and query speeds over a universal codecs for all data. As shown in **Figure 4**, different codec methods offer better or worse performance over different data types. Furthermore, column based compression allows for sophisticated filtering of records during decompression. Unlike tabix which must decompress all columns to check if a hit occurs (e.g., a p-value in some range), STABIX only needs to decompress a single column, reducing the time spent on data filtering and genomic position-based boundary checking. This feature allows for STABIX to improve on tabix significantly when there are no p-value hits present in a block, or when the p-value hits do not fall within the boundaries of a query. In these cases, STABIX can exit early and avoid unnecessary decompression while tabix queries necessarily still perform decompression before filtering (e.g. see GWAS file trait: size wine glass drunk). This phenomenon is demonstrated in **Figure 3C**, where STABIX decompression times for few p-value hits (e.g. 0-2) are much shorter than the same queries for tabix. **Figure 3D** shows a similar phenomenon where even when p-value hits are found in a block, if they occur outside of genomic position query boundaries STABIX can exit during the second filtering step and avoid full block decompression while tabix must decompress the entire block before p-value filtering.

STABIX’s configuration file allows for tunable parameters (e.g. block size and codec ensembles) to meet varying needs for different tasks (e.g. extreme compression for small file sizes or quicker decompression for frequent or many queries). We provide some baseline performance for 40 configures of 5 block sizes and 8 codec ensembles in **Figure 5, Table 2**, and **Supplementary Figure 1**. While we report these baseline performances for a single file and a p-value-based statistical index, we expect similar results for files of varying size and other columns of interest on which one might want to construct the index.. Depending on the expected number of hits, number of columns in the file, data types of those columns, and type of queries, different configurations can optimize for different performance goals. For example, on the same ten files, STABIX ensemble-xzb achieves a compression ratio of 1.13, while STABIX ensemble-xbb compression ratio is 1.4 (See **Supplementary Table 2)**.

## Supplementary Material, Tables, and Figures

**Supplementary Table 1.** Codec Ensemble Descriptions.

| Codec Ensemble Name | Codec used on Integers | Codec used on Floating Points | Codec used on strings |
| --- | --- | --- | --- |
| bz2 | bz2 | bz2 | bz2 |
| deflate | deflate | deflate | deflate |
| xz | xz | xz | xz |
| zlib | zlib | zlib | zlib |
| zstd | zstd | zstd | zstd |
| ensemble-fbb | FastpForVB | bz2 | bz2 |
| ensemble-xbb | xz | bz2 | bz2 |
| ensemble-xzb | xz | xlib | bz2 |

**Supplementary Table 2.** Performance of STABIX. Performance measurements for 10 per-phenotype files from PanUKBB. STABIX compression using default block size 2000 and codec ensemble xbb. Compression ratios include index files when available.

| trait | shellfish intake | sickle cell anemia | psoriasis | size of red wine glass drunk | standing height | monocyte count | type 2 diabetes | diabetes mellitus | snoring | smoking status |
| --- | --- | --- | --- | --- | --- | --- | --- | --- | --- | --- |
| no. sig genes | 977 | 42 | 143 | 0 | 5776 | 1803 | 141 | 156 | 59 | 168 |
| no. sig SNVs | 2189 | 392 | 2232 | 0 | 165522 | 41759 | 3514 | 4039 | 2120 | 5004 |
| avg. STABIX speedup over tabix | 0.7142 | 6.9028 | 7.1017 | 7.2302 | 4.0029 | 6.2065 | 7.6475 | 7.3949 | 7.6699 | 8.1163 |
| % STABIX wins | 0.2502 | 99.9896 | 99.9583 | 100.0 | 92.7063 | 97.8155 | 99.781 | 99.6768 | 99.8697 | 99.8488 |
| uncompressed (GB) | 1.2091 | 1.8225 | 4.1587 | 4.17 | 8.0126 | 8.7106 | 9.3567 | 9.3676 | 10.4422 | 10.6391 |
| bgzip (GB) | 0.2715 | 0.5107 | 1.0985 | 1.0982 | 2.0483 | 2.2893 | 2.4672 | 2.4732 | 2.7342 | 2.7522 |
| tabix index(MB) | 1.6885 | 1.7465 | 1.9274 | 1.9279 | 2.1941 | 2.2362 | 2.2698 | 2.2704 | 2.3161 | 2.3221 |
| STABIX (GB) | 0.1838 | 0.3333 | 0.7755 | 0.7788 | 1.5146 | 1.6266 | 1.7891 | 1.7933 | 1.9822 | 2.0288 |
| STABIX gen. index (MB) | 0.506 | 0.51 | 0.5127 | 0.5127 | 0.5186 | 0.5193 | 0.5202 | 0.5203 | 0.5211 | 0.5213 |
| STABIX stat. index (MB) | 0.1981 | 0.1534 | 0.1553 | 0.1537 | 0.2522 | 0.2024 | 0.1605 | 0.1604 | 0.1613 | 0.1698 |
| full STABIX comp. ratio | 6.5534 | 5.4573 | 5.3578 | 5.3498 | 5.2876 | 5.3527 | 5.2279 | 5.2217 | 5.2662 | 5.2423 |
| STABIX:bgzip+tabix comp ratio | 1.4805 | 1.5346 | 1.4177 | 1.4114 | 1.3531 | 1.4081 | 1.3798 | 1.3799 | 1.3801 | 1.3572 |

**Supplementary Figure 1.**
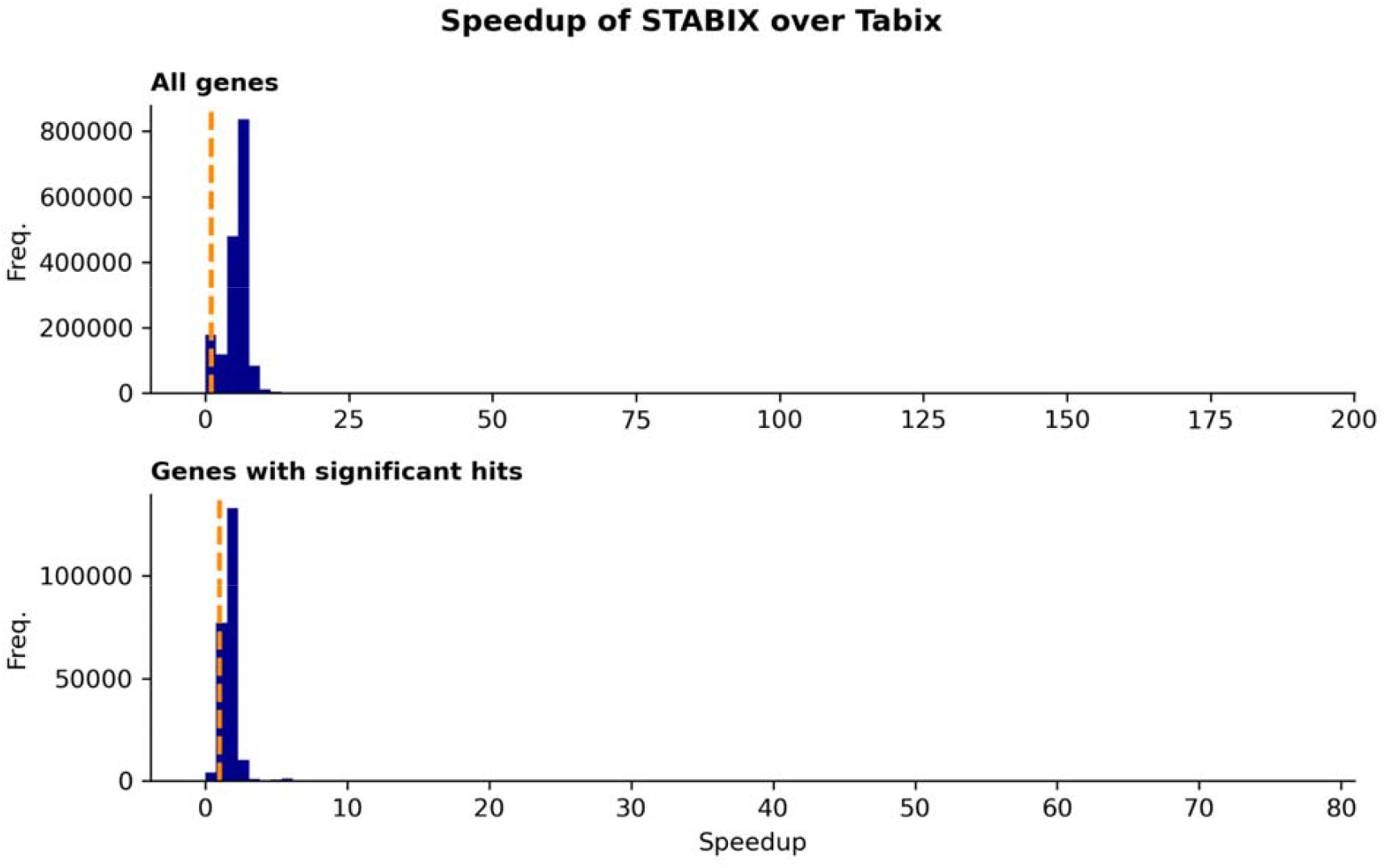
STABIX vs. tabix for 100 PanUKBB continuous files. Speedup by gene for significance SNV queries for STABIX over tabix + filtering step for all genes (top) and only genes with significant hits (bottom). A vertical dashed line (orange) is drawn at x=1 to separate where tabix wins (left of line) and STABIX wins (right of line). All figures report data for 100 per-phenotype files from PanUKBB, STABIX compression using default block size 2000 and codec ensemble xzb.

**Supplementary Figure 2.**
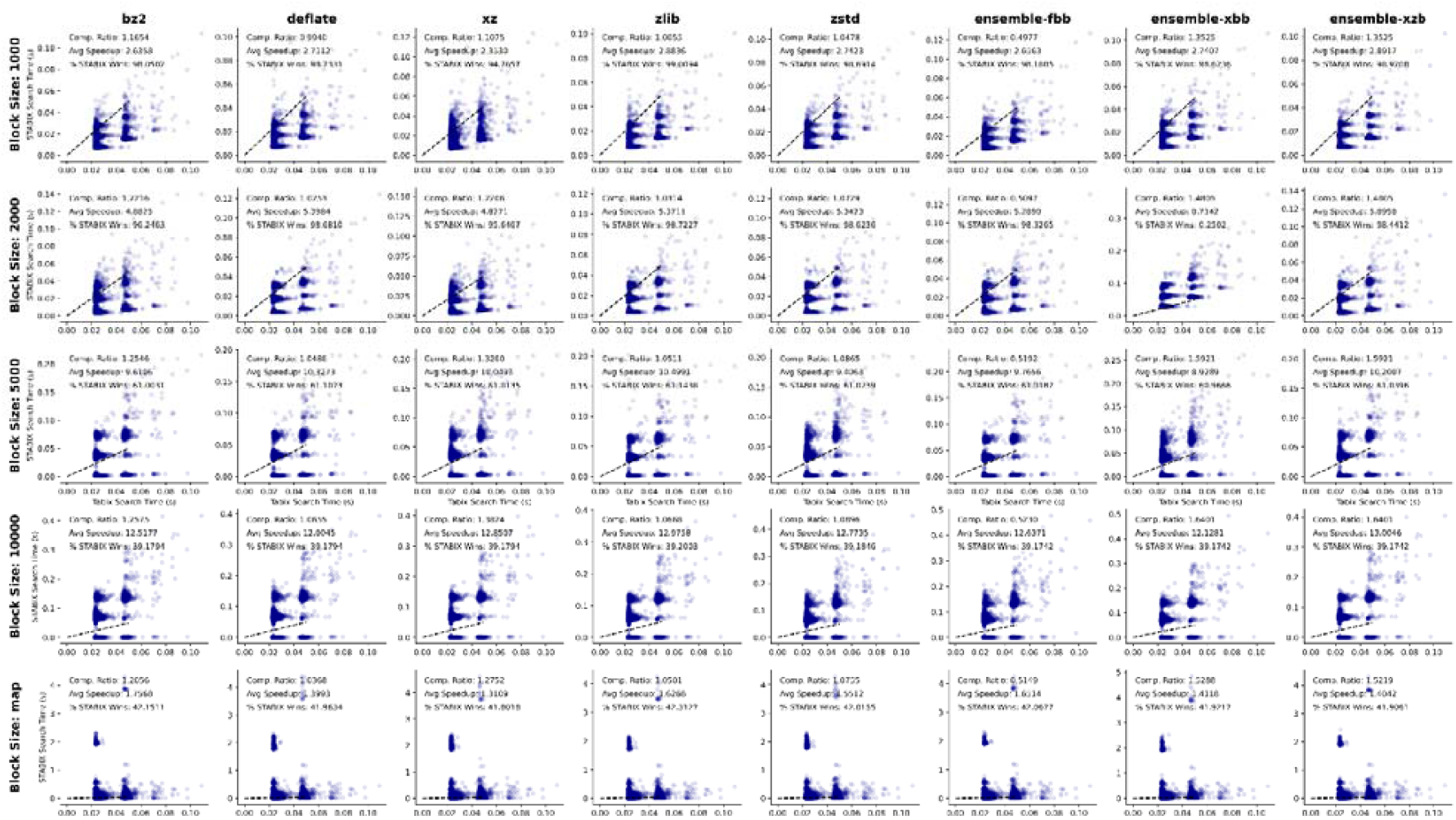
Complete combinations for 40 block size x codec ensembles. STABIX vs. tabix gene-based query times for PanUKBB GWAS file trait: shellfish intake. Times are shown for eight codec configurations from left to right: bz2, deflate, xz, zlib, zstd, ensemble-fbb, ensemble-xbb, and ensemble-xzb; and across five block sizes from top to bottom: 1000, 2000, 5000, 10000, and 1cM.. Bgzip + tabix : STABIX + genomic index + statistical index compression ratios, STABIX average speedup, and % STABIX wins are reported for each of the 40 configurations. A dashed linear line separates STABIX wins from tabix wins.

**Supplementary Figure 3.**
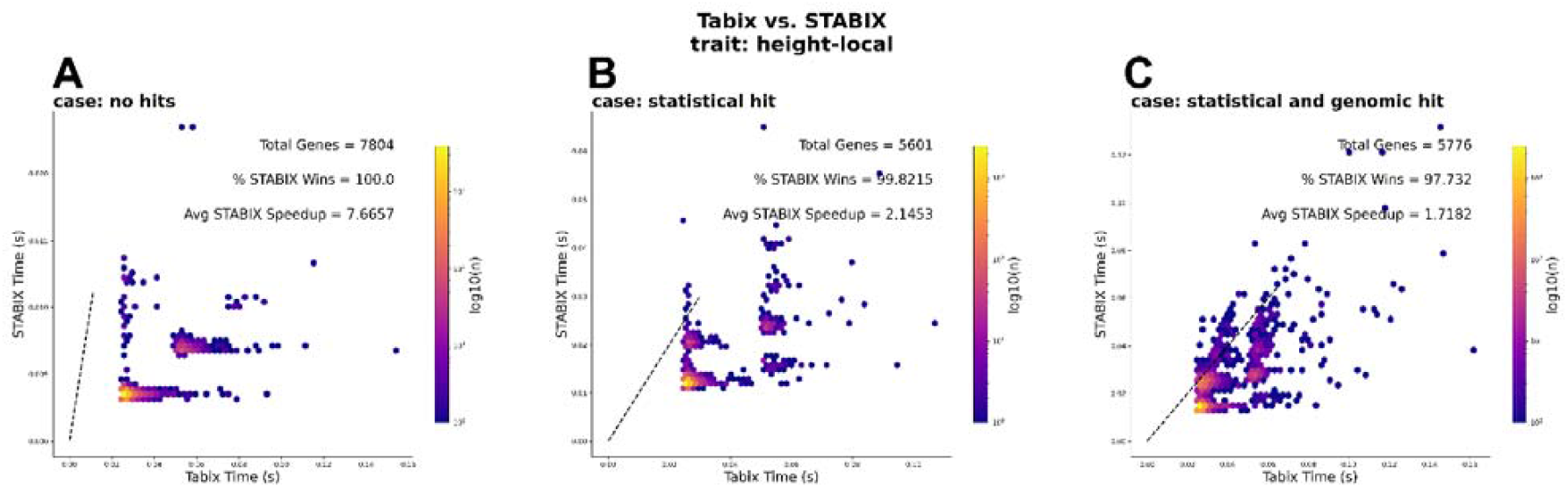
STABIX vs tabix search times by query case for standing height. STABIX vs. tabix gene-based query times for PanUKBB GWAS file trait: standing height. Each hexagonal bin is colored by the log of the number of genes included in that bin. **A** shows genes where there are no significant SNVs, **B** shows genes where there are significant p-value hits within a statistical query, and **C** shows genes where there are significant p-value hits within a statistical query and a genomic query. All plots include a dashed linear line which separates STABIX wins (below the dashed line) and tabix wins (above the dashed line).

### Author Credit. Kristen Schneider

–methodology and software development, data analysis and experimentation, manuscript writing. **Simon Walker**–software development. **Chris Gignoux**. Analysis support and supervision, manuscript review and editing. **Ryan Layer**–project supervision, methodology, manuscript review and editing.

### Data and Code availability

Software available for download at GitHub: https://github.com/kristen-schneider/stabix/.

Figures and analysis scripts available at GitHub: https://github.com/kristen-schneider/stabix-analysis

### Ten PanUKBB GWAS files

https://pan.ukbb.broadinstitute.org/docs/per-phenotype-files/index.html

continuous-103220-both_sexes: shellfish intake

phecode-282.5-both_sexes’: ‘sickle cell anemia

categorical-20096-both_sexes-2: size of red wine glass drunk

phecode-696.4-both_sexes: psoriasis

continuous-50-both_sexes-irnt: standing height continuous-30130-both_sexes-irnt: monocyte count

phecode-250.2-both_sexes’: ‘type 2 diabetes

phecode-250-both_sexes: diabetes mellitus

categorical-1210-both_sexes-1210: snoring

categorical-20116-both_sexes-0’: smoking status

### Map files

https://bochet.gcc.biostat.washington.edu/beagle/genetic_maps/

### Protein-coding genes BED file

Generated from the rom USCS table browser, we achieved a collection of 19,181 genes after collapsing regions of duplicate gene entries into a single region.

